# Targeting cancer-associated glycosylation for adoptive T cell therapy of gastro-intestinal and gynecological cancers

**DOI:** 10.1101/2024.09.12.612750

**Authors:** Andreas Zingg, Reto Ritschard, Helen Thut, Mélanie Buchi, Andreas Holbro, Anton Oseledchyk, Viola Heinzelmann, Andreas Buser, Mascha Binder, Alfred Zippelius, Natalia Rodrigues Mantuano, Matthias Matter, Heinz Läubli

## Abstract

CAR-T cell therapy has provided a significant improvement for patients with chemotherapy-resistant B cell malignancies. However, CAR-T cell treatment of patients with solid cancers has been more difficult, in part because of the heterogeneous expression of tumor-specific cell surface antigens. Here, we describe the generation of a fully human CAR targeting altered glycosylation in secretory epithelial cancers. The expression of the target antigen – the truncated, sialylated O-glycan sialyl-Thomsen-Nouveau antigen (STn) – was studied with a highly STn-specific antibody across various different tumor tissues. Strong expression was found in a high proportion of gastro-intestinal cancers including pancreatic cancers and in gynecological cancers, in particular ovarian and endometrial tumors. T cells expressing anti-STn CAR were tested *in vitro* and *in vivo.* Anti-STn CAR-T cells showed activity in mouse models as well as in assays with primary ovarian cancer samples. No significant toxicity was observed in mouse models, although some intraluminal expression of STn was found in gastro-intestinal mouse tissue. Taken together, this fully human anti-STn CAR construct shows promising activity in preclinical tumor models supporting its further evaluation in early clinical trials.

## Introduction

The treatment of advanced solid tumors by chemo- and immunotherapy remains challenging^1^. CAR T cells are genetically engineered T cells designed to recognize specific tumor antigens through chimeric antigen receptors and have revolutionized the treatment of hematological cancer^2,3^ but have not shown the same potency in solid cancers^4^. CAR-T cells are challenged with restricted access to tumor cells, an immunosuppressive tumor-microenvironment (TME), tumor-antigen heterogeneity and loss of tumor-antigens. Furthermore, antigen availability for CAR-T cell therapy is limited as solid tumor antigens are often expressed on healthy tissues at varying levels ^5^. Therefore, antigen selection is crucial in CAR design to ensure therapeutic efficacy while minimizing on-target off-tumor toxicity. These are often either specifically expressed on cancer cells or normal cells expressing it are deemed dispensable (e.g. CD19+ B-cells).

Differential gene-expression of enzymes involved in glycan synthesis and modification, including glucosidases, glycosyltransferases, and carbohydrate transporters, alters glycosylation patterns in cancer cells^6^. These alterations are often absent in normal cells, making them potential targets for directing CAR-T cells to the cancer microenvironment. Frequently found cancer-specific glycan modifications include increased branching and mannose content of N-glycans, increased sialylation and truncated O-glycans. Among these, the Sialyl-Thomson-Nouveau antigen (STn) was described as a cancer-specific marker^7,8^. It consists of a sialylated N-acetylgalactosamine (GalNAc) attached to a serine or threonine residue and is produced by the ST6GALNAC1 sialyl-transferase by adding sialic acid to the Tn antigen (Thomson-nouveau antigen, Supp. Data 1a)^9,10^. STn has been targeted therapeutically using various strategies, including radio-labeled antibodies^11,12^, cancer vaccines (Theratope)^13^ and first-generation CAR-T cells^14^. However, these approaches have not resulted in significant clinical benefit for cancer patients. Current pre-clinical research on glyco-targeted CAR-T cells focuses on glycosylated forms of Muc1^15,16^, with limited exploration of STn as a singular epitope^17^. Here, we constructed new fully humanized CAR-T cells targeting STn by using the highly specific 2G12-2B2 antibody as a backbone^18,19^.

## Results

### Immunohistochemical analysis shows increased expression of STn in neoplastic tissue

Tumor-restricted antigens that are frequently found in a large fraction of cancer patients are scarce. However, previous research indicated that STn might fulfill these requirements. Pan-tumor tissue microarrays (TMA) were stained with the 2G12-2B2 antibody (Fig. 1a) and revealed cancer types that showed frequent STn expression. Subsequently, we analyzed the ten cancer types with highest STn positivity along with healthy controls covering most tissues and calculated the area-based, pixelwise H-score^20^ (Fig. 1b, c). After careful review of tissue cores and their corresponding H-scores, we set the threshold for STn-positivity to H-score=10. Colon, uterine-cervical and pancreatic tumors resulted in the highest fractions (Fig. 1d) of STn-positive tumors which also applies to the median H-scores. Even though STn was found strongly overexpressed in many of the analyzed tumor types, its expression in healthy tissue was also observed, though at lower levels. Most STn expression was found in the digestive tract – from esophagus to colon – which is naturally lined with heavily glycosylated mucus (Supplementary Data 1c). Latest research suggests that STn modified mucus is crucial for its integrity and protection against excessive bacterial proteolytic degradation^21^. Small amounts of STn can also be found in the majority of analyzed tissue, such as pancreas, salivary glands, kidney and testis (Supplementary Data 1c).

**Figure 1.**
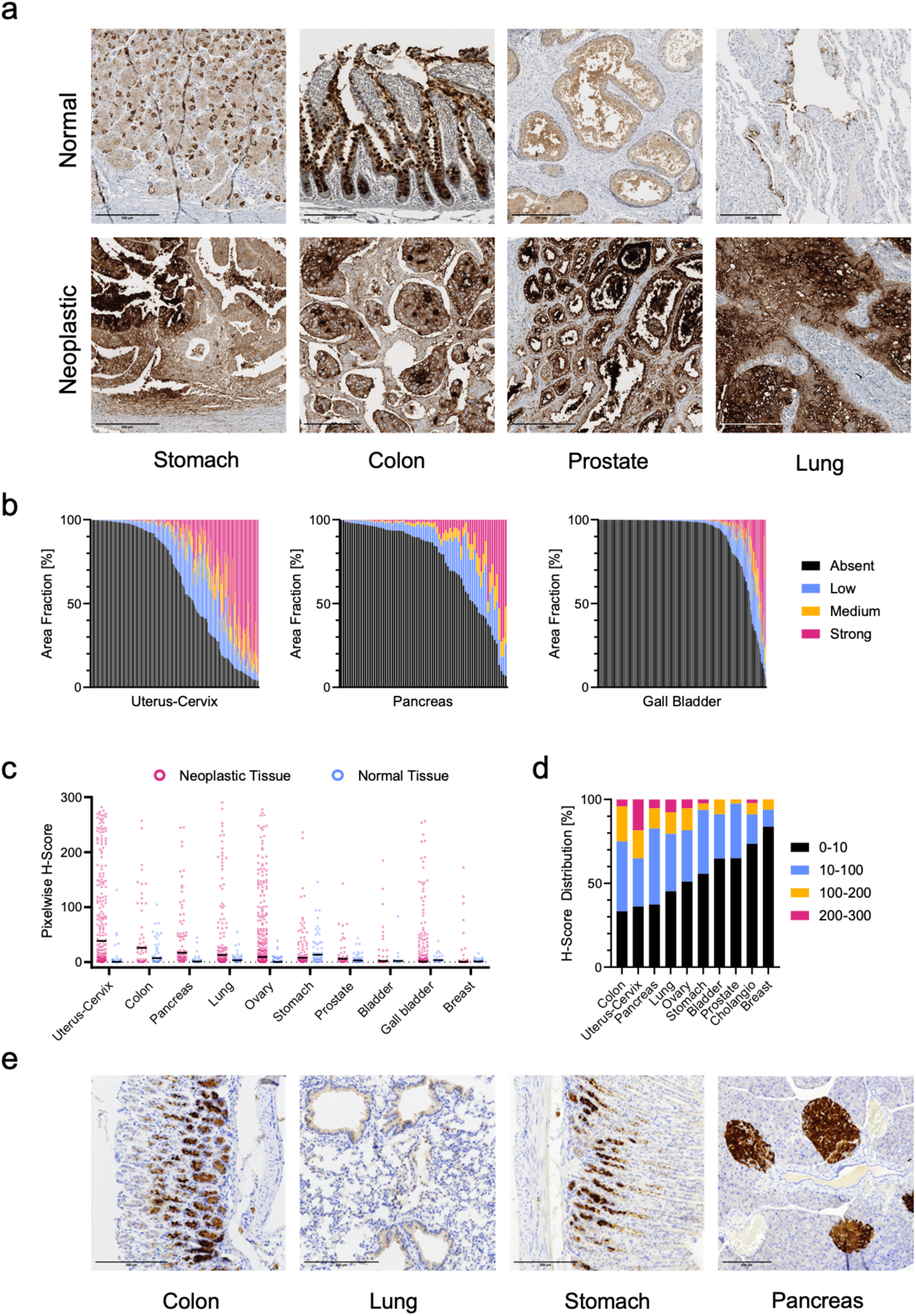
STn is overexpressed in neoplastic tissue of the gastrointestinal tract and female reproductive organs. **a**, Representative images of microarrays of healthy and neoplastic tissues stained for STn expression with the murine 2B2-2G12 antibody. Scale bar: 200 μm. **b**, Tissue cores were divided into negative, low, medium and high intensity staining by semi-automatic analysis in QuPath. Each column represents a TMA core from neoplastic tissue. **c**, The pixelwise H-score was calculated from the measured area-fractions. Each dot represents a single tissue core. Black bar: median. **d**, Pixelated H-scores were binned into 4 groups (H-score <10 was considered negative) and the fraction of each bin was calculated. **e**, STn expression in mouse organs. Positive staining detected in the colon, stomach and pancreas similar to human tissue. Some faint expression is visible also in the lung. Scale bar: 200 μm.

The area-based, pixelwise H-score itself is a valuable method for quantifying overall antigen expression in tissue samples. However, our analysis does not differentiate between intracellular and extracellular or membrane staining, which is critical for assessing the accessibility of the STn antigen for CAR-T cells and, consequently, their safety profile. In healthy tissue, STn expression is predominantly observed either intracellularly or on the luminal side as part of secretory products (Figure 1a, top row), which may reduce the risk of off-target effects. To predict potential toxicity in mouse models, we further tested STn-expression in mouse organs as glycans are often conserved (Fig. 1e). Positive staining was found in the colon, stomach and pancreatic islets. Faint STn expression was further detected in the lung epithelium. These four STn-positive tissues in mice corresponded to STn expressing tissues in human samples. Taken together, STn is highly expressed in cancers of the gastrointestinal tract, female reproductive system and lung and may therefore be a valuable target. Relevant toxicity of anti-STn CAR-T cells in humans cannot be excluded as relevant expression was observed in non-malignant tissues.

### The 2G12-2B2 scFv recognizes STn with high specificity and affinity

We generated a single-chain variable fragment (scFv) from the originally published humanized 2G12-2B2 sequence where the heavy and light chain is interconnected with a Whitlow-linker^22^. The scFv was fused to the IgG1-CH2-CH3 domains and recombinantly expressed. Despite the frequent expression of STn in solid tumors, few suitable cancer cell lines are available that stably express STn in cell culture conditions. Thus, we generated cell lines expressing the ST6GALNAC1^9^ sialyl-transferase enforcing expression of the STn epitope.

With these cell lines, we then verified the scFv-Fc’s specific binding and affinity to STn by flow-cytometry to cells that were positive or negative for STn expression. Transduced cell lines showed homogeneous expression of STn at different levels as shown by flow-cytometry when stained with the 2G12-2B2 scFv-Fc fusion protein (Fig. 2a). We further compared binding of the scFv-Fc protein with the full-length 2G12-2B2 antibody by titrating both on STn-positive and - negative A375 cells. The antibody had an affinity of 16 nM which is slightly higher than previously reported (K_D_ <10 nM) and did not show any binding to non-ST6GALNAC1-transduced cells. Surprisingly, the scFv-Fc titration resulted in an overall more intense signal and a lower K_D_ of ∼5 nM which might come from better epitope accessibility due to the smaller size. Also, no unspecific binding was observed for the scFv-Fc as well as for the antibody (Fig. 2b).

**Figure 2.**
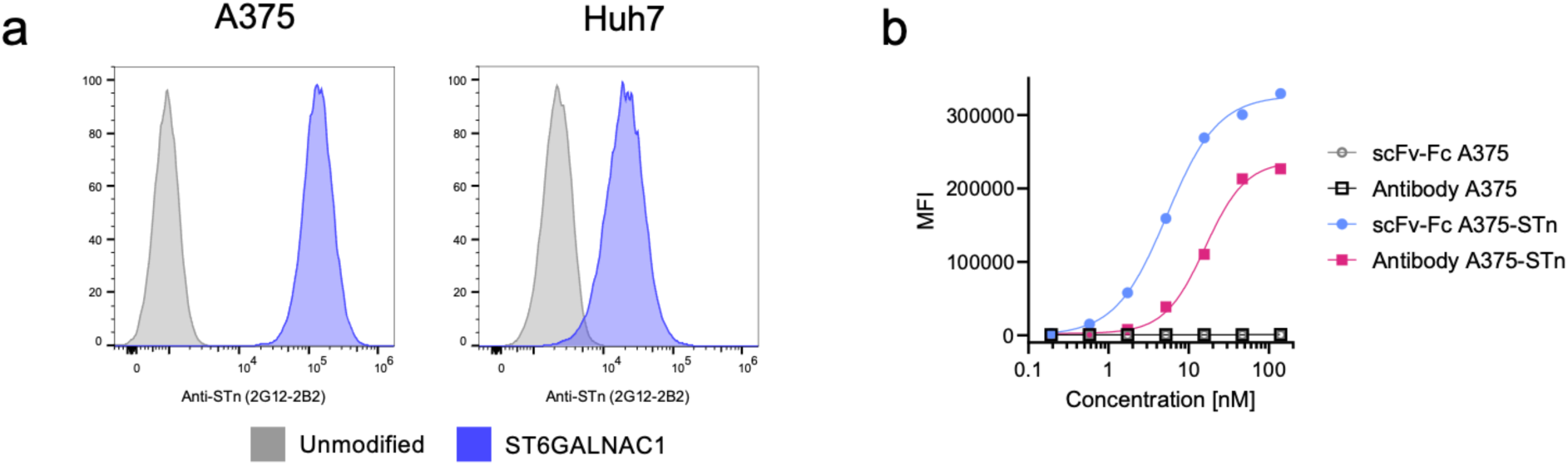
2G12-2B2 antibody and scFv-Fc bind tumor cell lines overexpressing ST6GALNAC1. **a**, Cancer cell lines stained with the anti-STn 2G2-2B12 scFv-Fc fusion protein. Overexpression of the ST6GALNAC1 sialyl-transferase results in STn expression. **b**, Affinity measurement of humanized scFv-Fc fusion protein compared to humanized 2G12-2B2 antibody on STn positive and negative cell lines. K_D_ antibody: 16 nM. K_D_ scFv-Fc: 5 nM.

### 2G12-2B2 anti-STn CAR-T cells are stably expressed and their phenotype is influenced by their signaling domains

The scFv was used in three different CAR configurations that have entered clinical trials or have been granted market authorization (Fig. 3a). The CD28-CD3ζ construct consists of the scFv, the extracellular, transmembrane and intracellular domain of CD28 followed by CD3ζ intracellular domain. The 4-1BB-CD3ζ has a similar configuration but the extracellular and transmembrane domains are derived from CD8 while the costimulatory domain is derived from CD137 (4-1BB). A rather new approach is followed with the TRuC (T-cell receptor fusion construct)^23–25^. Here, the scFv is directly fused to the full-length CD3ε protein via a (G_4_S)_3_ linker. We then transduced primary human T-cells and checked for CAR expression by staining with an anti-Whitlow-linker antibody^26^. Average transduction rates ranged from 50-90% and did not decrease over time. We consistently observed different expression levels of the CAR constructs. CD28-CD3ζ and TRuC constructs showed a medium expression level, while the 4-1BB-CD3ζ variant showed an increased expression level (Fig. 3b). We analyzed the phenotype of the CAR-T cells and classified them into stem-like memory (SCM: CCR7+CD62L+CD45RA+), central-memory (CM: CCR7+CD62L+CD45RO+) and effector/effector-memory (EFF-EM: CCR7/CD62L single and double negative) subtypes (Fig. 3c, Supplementary Data 2b). Few differences were observed for CD4 CAR-T cells and most of them (∼80%) acquired a more differentiated effector/effector-memory phenotype after two weeks of culture. In contrast, CD8 CAR-T cells more frequently retained the favorable stem-like memory phenotype. We observed a trend of 4-1BB-CD3ζ having a more differentiated phenotype which may have resulted from tonic signaling due to the higher expression level^27,28^. Excessive CAR expression was previously reported to be linked to early differentiation and exhaustion^27^. Further, previous reports clearly show that CAR-T products having a stem-like memory phenotype with short manufacturing times outperform those with a more differentiated phenotype or prolonged *ex vivo* culture^29,30^.

**Figure 3.**
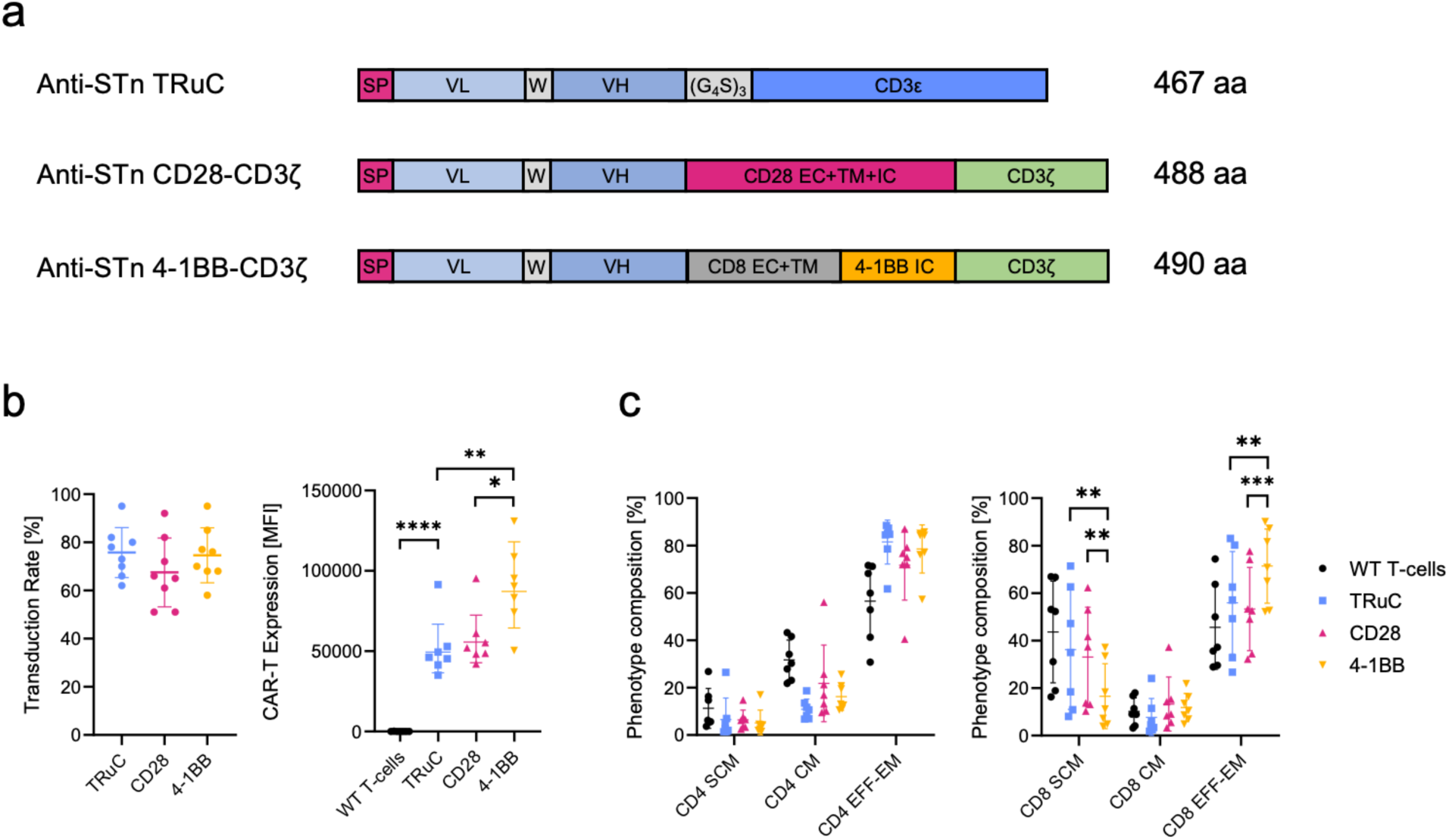
Anti-STn CAR is stably expressed on primary T-cells. **a,** Schematic overview of anti-STn CAR-T variants constructed using the 2G12-2B2 anti-STn scFv. SP: signal peptide, VL: variable domain light chain, VH: variable domain heavy chain, EC: extracellular domain, IC: intracellular domain, TM: transmembrane domain, W: Whitlow linker. **b,** Transduction rates and expression level of the CAR-T variants. Each dot represents CAR-T cells generated from PBMCs from a healthy donor. **c,** Phenotypic analysis of CAR-T cells. Stem-like memory (SCM): CCR7+CD62L+CD45RA+, Central memory (CM): CCR7+CD62L+CD45RO+, Effector/effector-memory (EFF-EM): CCR7 or CD62L single and double negative. WT: wildtype, untransduced T-cells. TRuC: anti-STn T-cell receptor fusion construct. CD28: anti-STn-CD28-CD3ζ CAR-T cells. 4-1BB: anti-STn-4-1BB-CD3ζ CAR-T cells. Comparison to untransduced T-cells not shown.

In brief, CAR-T cells that stably expressed the anti-STn chimeric antigen receptor were generated from a monoclonal antibody without reducing the affinity of the targeting scFv. Three different constructs were used that include two classic variants with CD28 or 4-1BB intracellular signaling domains as well the TCR-based *T-cell receptor fusion construct* TRuC.

### Anti-STn CAR-T cells show robust and prolonged cytotoxicity

Next, we wanted to test the cytotoxicity of engineered CAR-T cells. For this, we developed a cytotoxicity assay that allows quantifying the on- and off-target cell cytotoxicity, serial-killing capacity and *in vitro* persistence of different CAR-T constructs. We co-cultured the different CAR-T cells or untransduced T-cells at various effector-to-target (E:T) ratios with GFP-expressing target and non-target cell lines (Fig. 4a) and tracked the cytotoxicity by fluorescence-microscopy for five days. When tracking cytotoxicity over time at the E:T ratio 1:8, only small differences were observed for early timepoints between the CAR variants (Fig. 4b), especially for the Huh7-STn cell line. However, the cytotoxicity curves of the classic CAR-T cells started to separate from the untransduced T-cells and TRuC groups with prolonged co-culture. Both CD28-CD3ζ and 4-1BB-CD3ζ CAR-T cells showed sustained killing activity. The TRuC T-cells were not able to control the tumor cell growth at this E:T ratio. Despite some initial killing activity, mostly observed for the Huh7-STn cell line, the tumor cells eventually outpaced the TRuC T cells, growing faster than they could be eliminated. We hypothesize that the lack of additional co-stimulation leads to anergy due to inadequate T-cell signaling and despite mimicking native TCR engagement^31,32^.

**Figure 4.**
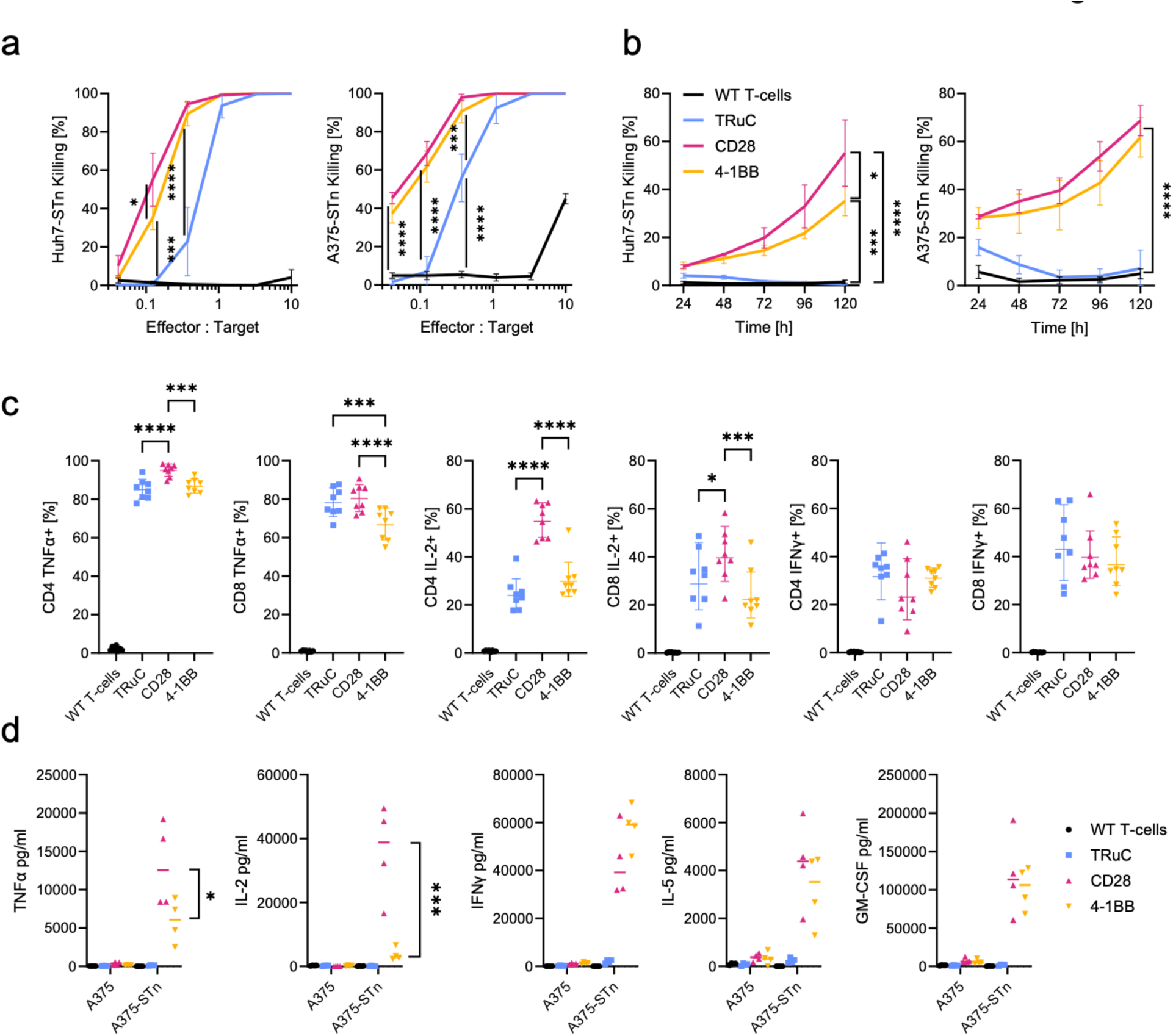
Classic anti-STn CAR-T cells show potent cytotoxicity. **a**, Cytotoxic activity of the anti-STn CAR variants assessed by co-cultures at different effector-to-target (E:T) ratios after 5 days of co-culture. **b,** Cytotoxic activity over 5 days. Effector-target-ratio = 1:8. Target cell lines: A375-STn and Huh7-STn. Pooled data from 4 donors. **c**, Intracellular cytokine staining normalized to CAR-expression for CD4 and CD8 CAR-T cells after 4 hours of co-culture with STn-expressing A375 cancer cells. Comparison to untransduced T-cells not shown. **d**, Analysis of secreted cytokines after 72 h of co-culture with A375 STn-negative or STn-positive cancer cells.

At elevated effector-to-target ratios, we observed notable off-target killing for all CAR constructs as well as for untransduced T-cells (Supplementary Data 3a). Of all CAR variants, TRuC T-cells showed the lowest degree of unspecific killing. After initial off-target cytotoxicity, the non-target tumor cells proliferated again up to full confluence or could at least balance the non-specific killing by T-cells, indicated by absence of killing compared to wells without effector cells (Supplementary Data 3c).

In a second setup, CAR-T cells from 4 PBMC donors were stimulated twice a week with an increasing number of target cells (Supplementary Data 4a) until target cells managed to adhere and start growing. The day after each stimulation cycle, activation and exhaustion markers were analyzed by flow cytometry. After the fourth stimulation cycle, tumor cells started growing in all TRuC co-cultures. Flow cytometry analysis showed a rapid decrease in 4-1BB expression and upregulation of PD-1 (Supplementary Data 4b). Co-cultures of CD28-CD3ζ and 4-1BB-CD3ζ CAR-T cells were terminated after 7 rounds of stimulation when tumor cells started adhering for most donors. Inability of completely clearing tumor cells was rather dependent on the donor than on the costimulatory domain (data not shown). Upregulation of 4-1BB was stronger on CD28-CD3ζ than on 4-1BB-CD3ζ CAR-T cells but decreased with time. Similarly, PD-1 was expressed constantly and stronger on CD28-CD3ζ CAR-T cells, an observation that has been reported previously^33,34^.

In summary, while the TRuC variant can specifically kill antigen-positive target cells and shows less off-target toxicity, it is not able to exert prolonged cytotoxic activity. In contrast, classic CAR-T cells with CD28 and 4-1BB costimulatory domains keep their cytotoxic activity even at very low E:T ratios.

### CD28-CD3ζeta CAR-T cells are potent cytokine producers

We then wanted to test the cytokine production of different engineered anti-STn-CAR-T cells. Cytokine production upon target-cell engagement is a crucial function of T-cells. Cytokines act in an autocrine manner on the T-cells themselves and in a paracrine manner on the tumor microenvironment thereby attracting other immune cells. We co-cultured the different CAR constructs with target and non-target cells and analyzed the cytokine production intracellularly (short-term) and in the supernatant (long-term). This allowed us to determine the immediate frequency of cytokine-expressing CAR-T cells as well as to compare the prolonged cytokine production by the different constructs. All variants were able to specifically induce TNFα, IFNγ and IL-2 production shortly after engagement with target cells (Fig. 4c). The most striking difference among CAR-T constructs was observed for intracellular IL-2 with significantly higher expression in CD4 CD28-CD3ζ CAR-T cells. Results from supernatant analyzed after 72 hours of co-culture, however, showed substantial differences between the classic and TCR-based CAR-T cells (Fig. 3d). The TRuC variant was unable to secrete cytokines for a prolonged time and only resulted in a minimal increase of measured cytokines in the supernatant. Contrastingly, the classic CAR-T cells with CD28 and 4-1BB costimulatory domains secreted high amounts of cytokines and mostly, though more pronounced, underline the data of intracellular cytokine staining. The most apparent difference is again found in IL-2 secretion.

The classic CAR-T constructs with CD28 or 4-1BB intracellular signaling domains showed some background expression in intracellular cytokine staining when co-cultured with non-target cells (Supplementary data 5b). This kind of tonic signaling is well known and might be one of the drawbacks of conventional CAR-T cells, eventually causing T-cell exhaustion^35,36^. The TRuC T-cells showed similar off-target cytokine expression as untransduced T-cells and thus indicate decreased tonic signaling.

Summarizing, we have compared short- and mid-term cytokine expression where we found striking differences between conventional and TCR-based CAR designs lacking costimulatory domain, but only during prolonged co-culture. CD28-CD3ζ and 4-1BB-CD3ζ CAR-T cells differed most in IL-2.

### Anti-STn CD28-CD3ζ CAR-T cells show potent in vivo tumor control without on-target off-tumor toxicity

For further investigation on the potency of the different anti-STn CAR-T cells, a subcutaneous tumor model in NSG mice was utilized, using the same cell lines as for the *in vitro* experiments. A375-STn melanoma were injected subcutaneously and grown for 15 days until they reached an average of 45 mm³. A single dose of 5 mio CAR-T cells or an equivalent number of untransduced T-cells were injected intravenously. No signs of immediate toxicity were observed regarding weight and body-score. Only the treatment of mice with CD28-CD3ζ CAR-T cells resulted in some cures and long-term tumor growth control (Fig. 5a).

**Figure 5.**
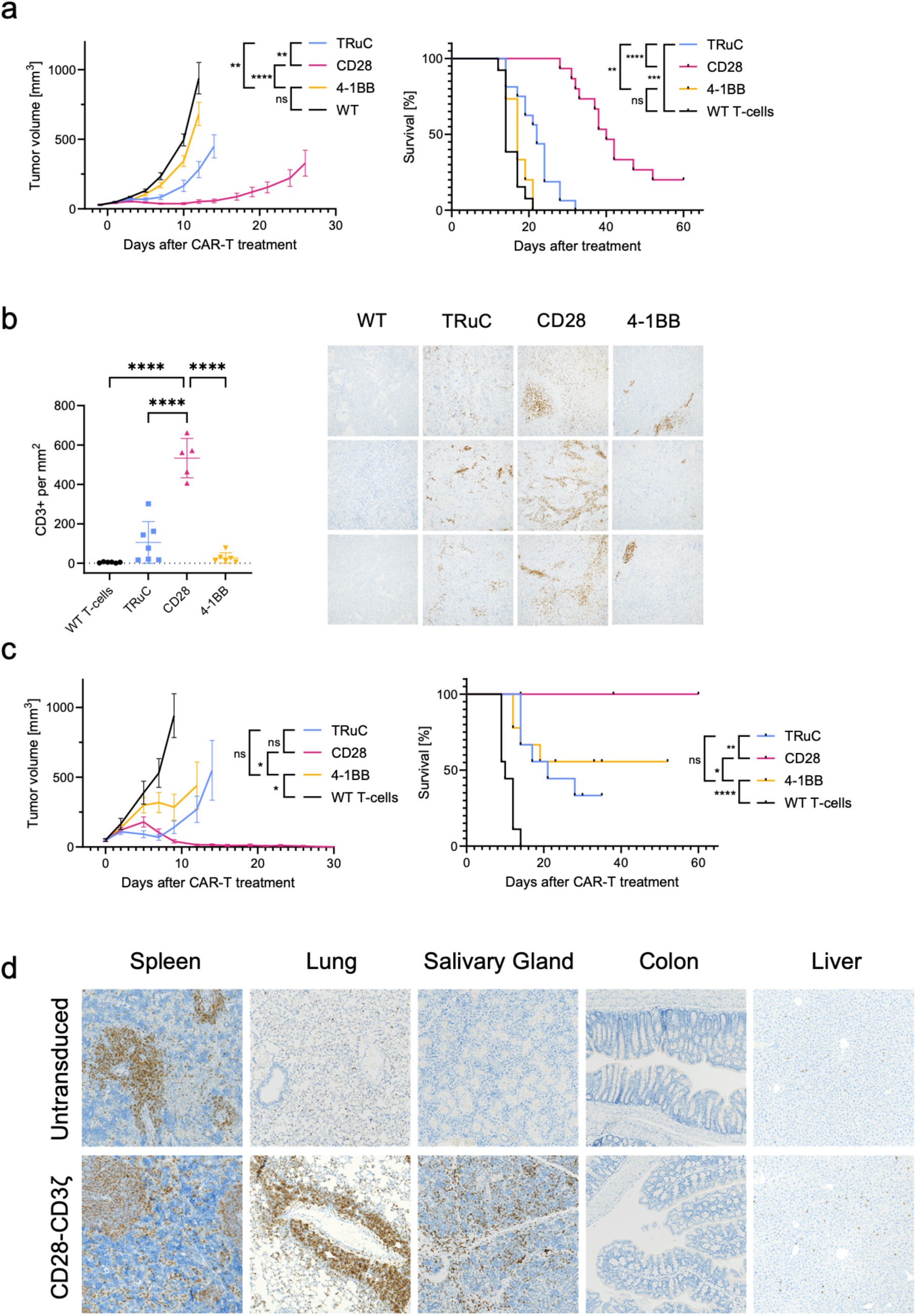
Anti-STn CAR-T cells with CD28 costimulatory domain effectively control tumor growth. **a,** Mean tumor volume with SEM and survival of NSG mice inoculated with A375-STn melanoma cells. Treatment of animals with 5 mio CAR-T cells when tumor volumes reached ∼45 mm^3^ (15 days). Pooled data of two independently performed experiments. Groupe size: 7-8 animals per experiment. **b,** CD3 infiltration analysis of A375-STn endpoint tumors. CD3+ cells per mm^2^ determined by QuPath analysis. **c,** Mean tumor volume with SEM and survival of NSG mice inoculated with Huh7-STn liver carcinoma cells. Treatment of animals with 15 mio CAR-T cells when tumor volumes reached ∼45 mm^3^ (9 days). Pooled data of two independently performed experiments. Groupe size: 4-5 animals per experiment. **d,** CD3+ infiltration analysis of normal mouse tissue by anti-STn-CD3ζ CAR-T cells and untransduced T-cells.

Tumors were harvested at endpoint and stained for infiltrating CD3^+^ cells. The groups of untransduced T-cells (WT: wildtype) and 4-1BB-CD3ζ CAR-T showed very little or only localized infiltration while CD28-CD3ζ CAR-T and TRuC treated animals showed a more abundant and distributed T-cell infiltration pattern (Fig. 5b). This was enough to slightly delay tumor growth for TRuC T-cells but was insufficient to completely cure tumors. The prolonged survival and tumor control of the CD28-CD3ζ CAR-T cells strongly correlated with infiltration density.

In a second tumor model to confirm previous findings, Huh7-STn hepatocellular carcinoma cells were injected subcutaneously in NSG mice. These tumors largely differ from the A375-STn model: while A375-STn tumor cells form a solid, whitish tumor, Huh7-STn tumors are soft and surrounded by blood. Access of CAR-T cells should therefore be less restricted.

9 days after tumor inoculation, 15 mio CAR-T cells or an equivalent number of untransduced T-cells were injected intravenously. In this model, CD28-CD3ζ CAR-T cells showed again superior tumor control and survival compared to 4-1BB and TRuC constructs. In this experiment, the 4-1BB construct showed a much better performance than in the A375-STn model, most likely due to the less restricted tumor access (Fig. 5c). The results closely recapitulate the outcome of the co-culture experiments in which the CD28-CD3ζ showed the best performance as well.

No significant off-tumor toxicity that resulted in severe weight-loss or reduced body score was noted. When animals were administered 15 mio CD28-CD3ζ CAR-T cells, we observed a subtle reduction in weight of which the animals timely recovered and no decrease in body score was observed (Supplementary Data 6a).

In a separate biodistribution experiment, male and female NSG mice were injected with 10 mio of the anti-STn CD28-CD3ζ CAR-T cells or an equivalent number of wildtype T-cells from two different PBMC donors. Mice did not bear tumors to avoid tumor-related expansion of CAR-T cells. 12 days later, mice were sacrificed and organs were harvested for human CD3 infiltration analysis (Fig. 5d, Supplementary Data 6b).

Unsurprisingly, CD3^+^ cells were abundantly found in the spleen for CAR-T and unmodified T-cell. When comparing organs from mice injected with anti-STn CAR-T cells to mice injected with unmodified T cells, we found increased infiltration of the lung and the salivary gland by CD3^+^ cells. In the lung, CD3^+^ cells accumulated mostly around the bronchia while in the salivary gland no specific pattern was found. The infiltration of the lung was not completely unexpected as migration of CAR-T cells towards the lung right after adoptive transfer was reported before in an antigen-independent manner^37^. Despite STn expression in the gastrointestinal tract, there was no increased T-cell infiltration after treatment with anti-STn CAR-T.

These animal experiments showed the superior *in vivo* efficacy of anti-STn CD28-CD3ζ CAR-T cells and good tolerability.

### Anti-STn CAR-T cells recognize natively expressed STn on patient-derived ovarian carcinoma cells

Previous experiments used cell lines strongly overexpressing STn. As the CAR/antigen interaction requires a higher antigen density for activation compared to classic TCR signaling^38,39^, we wanted to evaluate if our CAR-T cells can recognize native STn levels on patient-derived cancer cells.

We chose tumor digests of ovarian carcinoma as this cancer type showed frequent STn expression. Melanoma samples were used as negative controls as this tumor type is rarely positive for STn. Tumor digests were evaluated by flow cytometry for STn expression (Fig. 6a, Supplementary Data 7a). Additionally, STn-positive and negative cell lines were included as assay controls. We co-cultured the tumor cells with CAR-T cells generated from four healthy PBMC donors and analyzed their performance in multiple assays.

**Figure 6.**
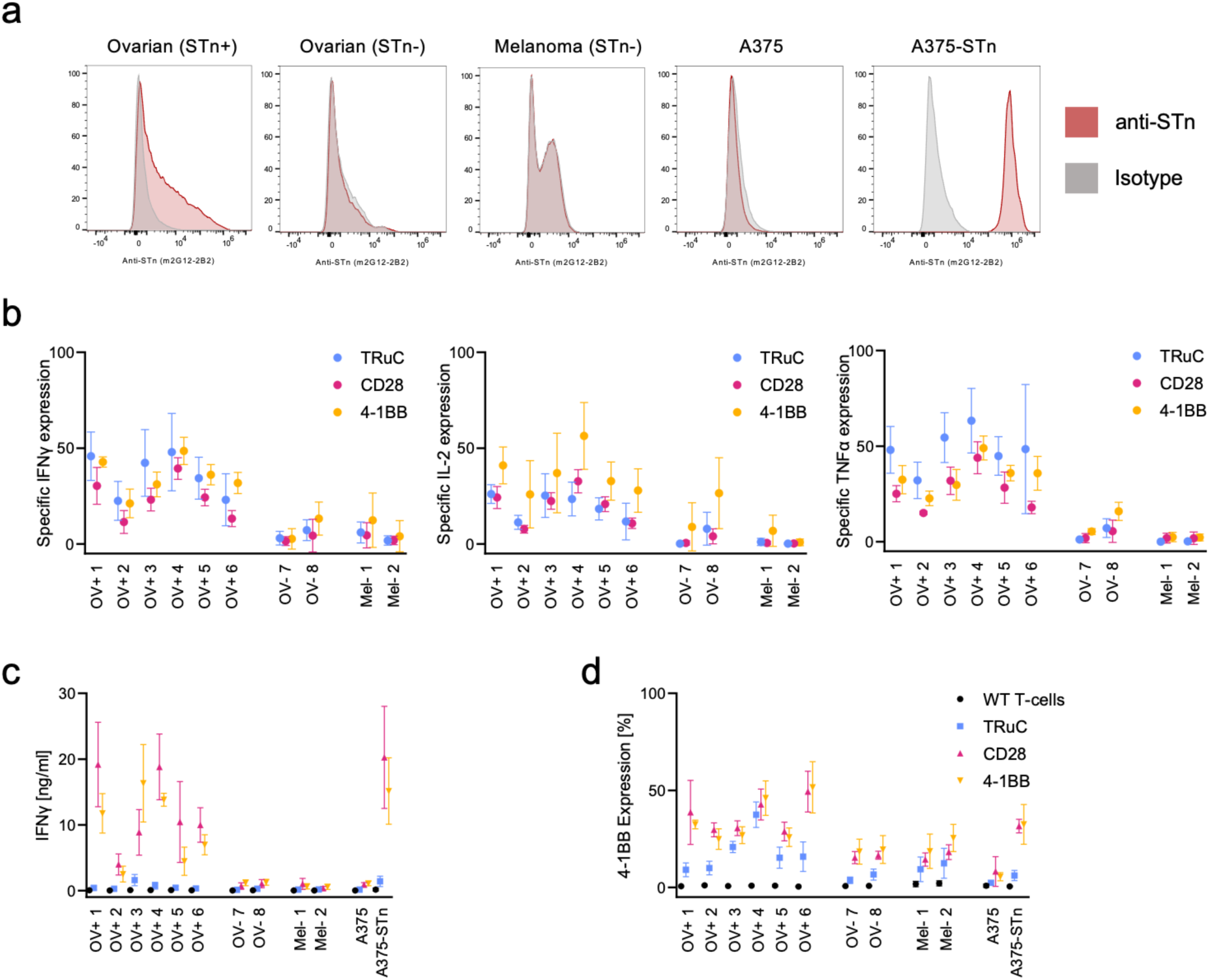
Anti-STn CAR-T cells recognize native STn-expression on primary ovarian carcinoma cells. **a,** STn expression on primary ovarian carcinoma patient samples and reference cell lines. Murine anti-STn antibody compared to isotype control, gated on DAPI-/CD45-. **b,** Intracellular cytokine staining of CD4+ T-cells after co-culture with tumor samples. Normalized to cytokine expression when cultured with STn negative and positive cell lines. OV+: Ovarian STn-positive, OV-: Ovarian STn-negative, Mel-: Melanoma STn-negative. **c-d,** IFNγ secretion and 4-1BB expression of CAR-T cells after co-culture with primary tumor samples or cell lines, normalized to CAR expression

Intracellular cytokine staining of IL-2, IFNγ and TNFα was performed after 6 hours of co-culture to evaluate antigen-specific activation of CAR-T cells (Fig. 6b, Supplementary Data 7b). For better comparability of antigen-specific induction of cytokine expression, we normalized the frequency of cytokine-expressing cells to the expression of the control cell lines. All CAR variants selectively started producing cytokines upon engagement with STn-positive tumor cells but not or only very weakly on STn-negative cells and no considerable differences were observed. Unmodified T-cells did not show any cytokine expression above background.

Further, after three days of co-culture, upregulation of 4-1BB and IFNγ secretion was analyzed (Fig. 5c, d). In sharp contrast to the intracellular cytokine staining, TRuC CAR-T cells failed to secrete notable amounts of IFNγ compared to classic CAR-T cells, independent if co-cultured with primary tumor cells or cell lines as observed before. A similar pattern can be seen with 4-1BB upregulation. TRuC CAR-T cells showed a lower level of 4-1BB upregulation.

Interestingly, upregulation of 4-1BB on CAR-T cells was higher in certain tumor samples than with the positive control cell line.

These results show that our CAR-T cells can recognize STn on primary tumor material while ignoring antigen-negative cells. Best results regarding persistent T-cell activation and IFNγ secretion was observed for classic CAR-T cells which is in line with results observed from *in vitro* killing assays and mouse models.

## Discussion

The engineering of CAR-T cells to treat solid cancer is very challenging. Here, we evaluated the Sialyl-Thomson-Nouveau (STn) antigen as a CAR-T tumor target using the recently developed, humanized anti-STn antibody 2G12-2B2. Staining of various tumor entities revealed frequent and strong STn expression of tumor cells, especially in those that have strong secretory activity such as uterine-cervical, colon, pancreatic and lung cancer. Most of the STn-staining in healthy tissue was observed intracellularly or on the luminal side of the gastrointestinal tract. Still, the degree of STn on the cell surface of healthy cells and its accessibility for CAR-T cells remains unclear. The question of safety and tolerability in humans can thus not be fully answered. Certain quantities of antigen might be tolerated in non-inflamed tissues as shown in clinical trials using B7H3-, EpCAM- or PSCA-targeted CAR-T cells ^40–42^. Few mouse organs such as stomach and colon also showed a similar expression pattern as in human tissue.

We developed, characterized and tested multiple chimeric-antigen-receptor constructs using well known conventional configurations with CD28 or 4-1BB and CD3ζ intracellular domains as well as the TCR-integrating TRuC variant. *In vitro* cytotoxicity assays revealed differences over time between the classic and TCR-based constructs. CD28 and 4-1BB based CAR-T cells were able to control the growth of tumor cells at lower E:T ratios than TRuC T-cells. Cytokine secretion was high for CD3ζ-based CAR-T cells but low for the TRuC. Anti-STn-CD28-CD3ζ showed the strongest secretion of TNFα and IL-2. We hypothesize that the lack of co-stimulation (Signal 2) in the TRuC causes incomplete T-cell signaling and activation, ultimately leading to dysfunction and exhaustion. In repetitive stimulation and cytotoxicity assays, the TRuC T-cells were rapidly overgrown by tumor cells. Xenograft models partially confirmed these findings. TRuC T-cells showed some effect in the A375-STn mouse model but did not result in cures despite tumor infiltration. Our findings contrast previous reports on the TRuC construct. While others reported superiority of the TRuC in *in vivo* models ^23,25^, in context of STn-targeting CAR-T cells, the TRuC had inferior *in vivo* and *in vitro* efficacy. The anti-STn CD28-CD3ζeta CAR-T cells on the other side showed the best performance in most experiments. They produced high amounts of cytokines, show prolonged and potent *in vitro* killing activity and could best control the tumor growth in both mouse models. Further, treatment of mice with anti-STn CAR-T cells did not result in chronic or lethal toxicity, suggesting limited accessibility of the antigen for systemically applied anti-STn-CAR T cells.

*In vitro* analysis gave some hints why the 4-1BB-CD3ζ construct performed inferior in *in vivo* models, e.g. lower IL-2 secretion. However, it remains unclear why the tumor infiltration in the A375 mouse model is minimal compared to the other CAR-T groups. No similar findings were reported before. The optimal combination of scFv and CAR construct might also be target and context dependent as there is no consent yet – in preclinical as well as in clinical settings – which costimulatory domain is the best. For example, from eight preclinical studies targeting CD19 in B-cell malignancies with second-generation CAR-T cells, five reported CD28 to be the costimulatory molecule of choice, in two 4-1BB performed better while in one study there was no difference. Direct comparison of studies is especially complex as manufacturing, the overall architecture and the viral platform of the anti-CD19 CAR-T cells often differ as well as pre-treatment regimes^43,44^.

The sialyl-Thomson-Nouveau antigen has been previously known as a tumor marker but no therapeutic effects were reported when targeted by first-generation CAR-T cells, radio-labeled antibodies or cancer-vaccines. No severe adverse events were reported in context of STn-targeting. The development of novel, more specific, humanized anti-STn antibodies in recent years hopefully results in more effective future treatments that could affect many patients’ lives. Our STn-directed second-generation CD28-CD3ζ CAR-T cell might be another promising treatment strategy. No contraindications were identified that would immediately exclude STn as a promising CAR-T therapy target.

It is possible that our second-generation anti-STn directed CAR-T cells will have a reduced efficacy due to insufficient tumor infiltration in patients, T-cell exhaustion and reduced persistence as often observed in previous clinical trials^45^. There are several promising approaches for armored, fourth-generation CAR-T cells that address these challenges. Tumor homing can be improved by expressing chemokine receptors^46^, persistence can be increased by inducible or constitutive cytokine expression^47,48^ and exhaustion can be reduced by decoy or switch receptors inhibiting PD1^49^ or TGFβ^50^ signaling. Some approaches are already in clinical trials, ranging from promising results (e.g. NCT04684563, NCT05715606) to unfortunate and even fatal outcomes (NCT04227275) due to adverse effects.

As we have found a strong expression in tumor tissues of patients with epithelial ovarian cancer and also pancreatic cancer, a clinical trial in these patient populations with a high-medical need would be the logic next step. In a phase 1 clinical trial, the safety profile of anti-STn CD28-CD3ζ cells will be established. A trial using the *Sleeping Beauty* transposase system^51^ is therefore planned within our institution.

Taken together, we constructed a new, effective CAR that is targeting a glycan-tumor antigen. We found that anti-STn CAR-T cells can be successfully activated by primary gynecological cancer cells and effectively eliminates tumor in mouse models. Further testing in early clinical trials for this approach is needed.

## Materials and Methods

### Plasmid construction

All plasmids were planned *in silico* with Geneious Prime (Version 2024.0.3). DNA sequences were acquired from publications^18^, patents^52,53^, Addgene or the NCBI database. Protein-coding regions were codon-optimized with Genscript condon optimizer, ordered as DNA fragments from Genscript or Twist Bioscience and cloned into a lentiviral plasmid backbone (pLV, VectorBuilder) under the control of the EFS (EF1α-short) promoter.

If required, restriction sites were introduced by primers (Microsynth) during PCR amplification (Q5, New England Biolabs). DNA fragments and plasmids were digested using FastDigest enzymes (Thermofisher) and ligated with T4 ligase (Thermofisher). Ligations were mixed with in-house made Stbl3 competent cells^54^, incubated for 20 min on ice and directly plated on prewarmed LB-plates containing 100 ug/ml ampicillin (Sigma).

When restriction-cloning was not suitable, DNA fragments were assembled with the GeneArt Gibson Assembly HiFi Master Mix (Thermofisher). All plasmids were verified by colony PCR and sanger-sequencing (Microsynth).

### Cell Culture

A375-Luc2 were acquired from the American Type Culture Collection (ATCC) in which the β2m gene was knocked-out (kind gift from S. Tundo, Cancer Immunology Lab, University of Basel). The Huh7 cell line was a kind gift from F. Stenner (University Hospital Basel). Cell lines were grown in RPMI supplemented with 10% FBS, 1% penicillin/streptomycin, 1% non-essential amino acids and 0.25 ug/ml amphotericin B. Lenti-X HEK293T cells were acquired from TakaraBio and grown in DMEM containing 10% FBS, 1% penicillin/streptomycin and 0.25 ug/ml amphotericin B. Cell lines were regularly tested for mycoplasma contamination.

### Lentivirus production

16 hours before transfection, 14 mio Lenti-X 293T cells (TakaraBio) were seeded in 18 ml complete DMEM medium (Sigma) into 15-cm culture dishes. For the transfection-mix, 2.5 ug pMD2.G, 4.7 ug pCMVR8.74 and 7.2 ug pLV transfer vector were mixed in 1.8 ml jetOPTIMUS buffer (Polyplus). 18 ul jetOPTIMUS was added and incubated for 10 min before adding the transfection-mix to the cells. Medium was exchanged after 5-6 hours and lentiviral particles were collected after 24 and 48 hours after medium exchange. The pooled supernatant was concentrated with 4X in-house made PEG-8000 solution and resuspended in 1 ml Prime XV serum-free T-cell medium. Aliquots were stored at −80°C.

### Antibody and scFv-Fc recombinant protein expression

CHO-S cell were cultured in HyCell TransFx C-Medium (Cytiva) at <8 mio/ml on a shaking platform. At the day of transfection, cells were isolated and resuspended in fresh medium at 2 mio/ml. 6-8 hours after medium exchange, cells were transfected with FectoPRO according to manual (0.8 ug DNA and 1.2 ul FectoPRO per ml of total volume). 24 hours after transfection, culture was diluted twofold. 10 days after transfection, supernatant was harvested and antibody or scFv-Fc fusion protein was isolated with protein G resin, dialyzed against PBS and concentrated with Vivaspin centrifugal concentrators (Sartorius).

### Generation of CAR-T cells

Peripheral blood mononuclear cells (PBMCs) were isolated from buffy coats (Blutspendezentrum SRK beider Basel) by density centrifugation with Lymphoprep using SepMate tubes (Stemcell Technologies). Total PBMCs were then activated (10 mio PBMC/ml) with CD2/CD3/CD28 Immunocult T-cell activator (Stemcell Technologies) in Prime XV chemically-defined T-cell medium (IrvineScientific), supplemented with IL-15 (10 ng/ml, Peprotec) and 1% penicillin/streptomycin (Sigma). 2 days after activation, cells were isolated and resuspended in 2.5 ml medium containing 0.5 mg/ml synperonic F108 (Sigma) and 0.125 uM BX795 (MCE). 100 ul of concentrated lentivirus was added and cells were cultured in 6-well GRex plates. After 48 hours, wells were filled up with Prime XV supplemented with 10 ng/ml IL-15 and 1% penicillin/streptomycin and expanded for 14 days. For untransduced T-cells (WT T-cells), instead of lentivirus, Immunocult was added again but were treated otherwise the same. Medium was exchanged every 5-7 days. Experiments were performed within 2-3 weeks after production start.

### Generation of STn-expressing target cell lines

mRNA from OVCAR-3 cells was isolated, reverse-transcribed and amplified by PCR using primers specific for the ST6GALNAC1. DNA was cloned into a lentivirus transfer vector under control of an EFS promoter and lentivirus was produced as described. For cytotoxicity assays, target and non-target cell lines were further transduced with a lentivirus encoding the *green-fluorescent protein* GFP that allowed quantifying the area covered by cells of cell-culture plates via fluorescence-microscopy.

### Mouse models

NSG mice (NOD.Cg-Prkdcscid Il2rgtm1Wjl/SzJ) were bred in the animal facility of the Department of Biomedicine (University Hospital Basel) and experiments were performed under approval of the local Ethical Committee (Basel Stadt) and complied with ethical regulations. For *in vivo* tumor growth experiments, 3 mio A375-STn or 2 mio Huh7-STn cells were injected subcutaneously in the right flank. 9 days (Huh7-STn) / 15 days (A375-STn) after xenograft inoculation, when average tumor size reached ∼45 mm^3^, mice were injected intravenously with 15 mio (Huh7-STn) / 5 mio (A375-STn) CAR-T cells or an equivalent number of untransduced T-cells in PBS + 1% human serum albumin. Tumor size was measured three times a week with a caliper (0.5*width^2^*length) and mice were sacrificed before tumor volume exceeded 1500 mm^3^ or when significant discomfort was observed (e.g. eye infection, weight loss >15%).

For the biodistribution study, 8 male and 8 female non-tumor bearing NSG mice were injected intravenously with 10 mio CAR-T cells or an equivalent number of expanded wildtype T-cells. 12 days after injection, mice were sacrificed and organs were harvested.

Experiments were performed twice independently using different PBMC donors for CAR-T cells production.

### Flowcytometry

Cells were analyzed on CytoFLEX flow cytometers. All stainings were performed in PBS with 1% FBS and 2 mM EDTA with incubation steps at 4°C for 30 min. DAPI was used as a live-dead marker if not otherwise mentioned. CAR-T cells were stained with recombinantly expressed anti-Whitlow-linker antibody “Clone 16”^55^ (Biointron) and goat-anti-rabbit-PE antibody (Invitrogen). Cell lines were analyzed for STn expression by recombinantly expressed anti-STn scFv-Fc and anti-human IgG-AF647 (Biolegend).

### Cytokine analysis

CAR-T cell concentration was adjusted to 50% CAR-expression with untransduced T-cells. 100k T-cells were mixed with 50k A375 or A375-STn cells (E:T ratio 1:1) in complete medium and 5 ug/ml brefeldin A (Biolegend). Co-cultures were incubated for 3.5 hours in a cell culture incubator. Intracellular cytokines were stained with the Fix & Perm cell fixation & cell permeabilization kit (Thermofisher), using Zombie violet (Biolegend) as live/dead marker. To determine the MFI, gating included only cytokine-positive cells for CAR constructs while for untransduced T-cells the whole population was used for comparison.

The assay was performed twice in triplicates with T-cells from 4 different PBMC donors each. Cytokines in supernatant after 72 hours of co-culture were analyzed by ProcartaPlex Immunoassay (Thermofisher) according to manual.

### Cytotoxicity assay

10’000 GFP+ target or non-target cells were plated in 96-well flat bottom plates in complete RPMI medium without cytokines. 6 hours later, CAR-T cells were diluted to 50% CAR-expression with untransduced T-cells, 3-fold serially diluted and added to target cells, resulting in an effective 10:1 CAR-to-target-cell ratio for the highest dilution. An equivalent number of untransduced T-cells was added as control. The occupied area-fraction of GFP+ target cells was measured by fluorescence microscopy every 24 hours after addition of T-cells for 5 days. The assay was performed in duplicates/triplicates with T-cells from 4 different PBMC donors.

### Immunohistochemistry

Mouse organs were fixed in paraformaldehyde solution (37% stock solution diluted tenfold in PBS) for 48 hours and then paraffin-embedded on a TPC15 TRIO machine. Sections were processed on the Leica Bond system. In brief, antigen-retrieval was performed using citrate buffer (pH=6) for 30 min at 100°C. Humanized anti-STn scFv-Fc was used at 5 ug/ml for primary staining, followed by rabbit anti-human IgG (ab97156) and HRP/DAB polymer staining for detection. A monoclonal human IgG1 antibody was used as isotype control.

Human tissue microarrays were stained at the Institute of Pathology of the University Hospital Basel using the Ventana Benchmark Ultra system. The protocol optiView with pre-treatment CC1 for 16 min was followed. The murine 2G12-2B2 antibody (Genscript) was used at 3 ug/ml. Tissue microarrays (TMA) were analysed using QuPath.

Pixelwise H-Score^20^ was calculated from automated tissue and DAB detection by QuPath for unbiased quantification of the stained microarrays. TMAs were de-arrayed and tissue was detected by the classifier function for hematoxylin staining. Where necessary, tissue was manually excluded (tissue double-layers, out-of-focus areas etc.). Detected tissue was then classified by thresholding in no staining (<0.2), low (0.2-0.4), medium (0.4-0.6) and strong staining (>0.6). H-Score was calculated according to the following formula: H-Score = 100*(1*LOW_area + 2*MEDIUM_area + 3*STRONG_area) / (ZERO_area + LOW_area + MEDIUM_area + STRONG_area). Due to the heterogeneous structure and partially very intense staining of the tissue, it was not possible to computationally isolate the membrane staining only. Consequently, we used the weighted intensities of the areas instead, normalized to the total tissue area.

Mouse tumors for CD3 infiltration analysis were embedded in OCT after harvest. Cryosection were fixed with PFA solution. CD3 staining was performed on the Roche Ventana System using polyclonal rabbit anti-human CD3 antibody (Abcam, ab5690) by the Histology Core Facility (University of Basel).

### Co-culture of CAR-T cells and primary tumor samples

Tumor digests were stained for STn expression with murine 2G12-2B2 anti-STn antibody. STn staining was analyzed on DAPI/CD45 double-negative cells comparing to an isotype control. Assays were performed in AIM-V + 10% FBS + 1% penicillin/streptomycin. For intracellular cytokine staining, 5 ug/ml brefeldin A was included into the assay medium. Tumor digests or cell lines were labeled with cell-trace violet (Thermofisher, 1:4000 in 5 ml PBS, 30 min at room temperature). Labeled cells were equally distributed into assay wells in duplicate (100’000 live cells maximal per well) and 50’000 CAR-transduced T-cells or unmodified T-cells were added. After 6 hours of co-culture, intracellular cytokine staining was performed as outlined above. Supernatant was harvested after 3 days of co-culture and IFNγ secretion was analyzed using the Lumit human IFNγ Immunoassay (Promega) according to the manual and T-cells were analyzed for 4-1BB expression by flow cytometry.

### Statistics

GraphPad Prism version 10.1.2 was used for statistical analysis and for visualization of data. The Log-rank (Mantel-Cox) method was used to compare survival curves of mice treated with CAR-T cells. Tumor volume was compared at the timepoint where the first mouse had to be sacrificed. Tumor volumes were log-transformed after zeros were replaced with 1’s. Two-way ANOVA was performed followed by Tukey’s multiple comparisons test. Fluorescence intensities were log-transformed followed by one-way ANOVA with Tukey’s correction for multiple comparisons. If not otherwise stated, the mean and standard deviation is shown.

K_D_ was calculated by nonlinear curve fitting with variable slope (four parameters).

P-values were assigned as followed: P≤0.05 (*), P≤0.01 (**), P≤0.001: (***), P≤0.0001 (****).

## Contributions

A.Zin., N.M.R. and H.L. designed and planned the experiments. A.Zin. performed the experiments. M.Bu. helped with animal experiments. R.R. and H.T. are responsible for the transfer to the GMP production level. M.M. helped with the pathology analysis and A.B. and A.H. provided apheresis products. V.H. and A.O. provided clinical material and analysis. A.Zip. L.J., and M.Bi. provided relevant material. All authors read and approved the final manuscript.

## Acknowledgement

We thank Anna Katherina Stalder Cramm (Pathology University Hospital Basel) for developing and performing the STn staining on human tissue microarrays, Ewelina Bartoszek-Kandler (Department of Biomedicine Microscopy Core Facility) for helping with the bioinformatic analysis of human tissue microarrays and Diego Calabrese (Department of Biomedicine Histology Core Facility) for performing the CD3 staining of mouse tissues.

## Supplementary Data

**Supplementary Data 1.**
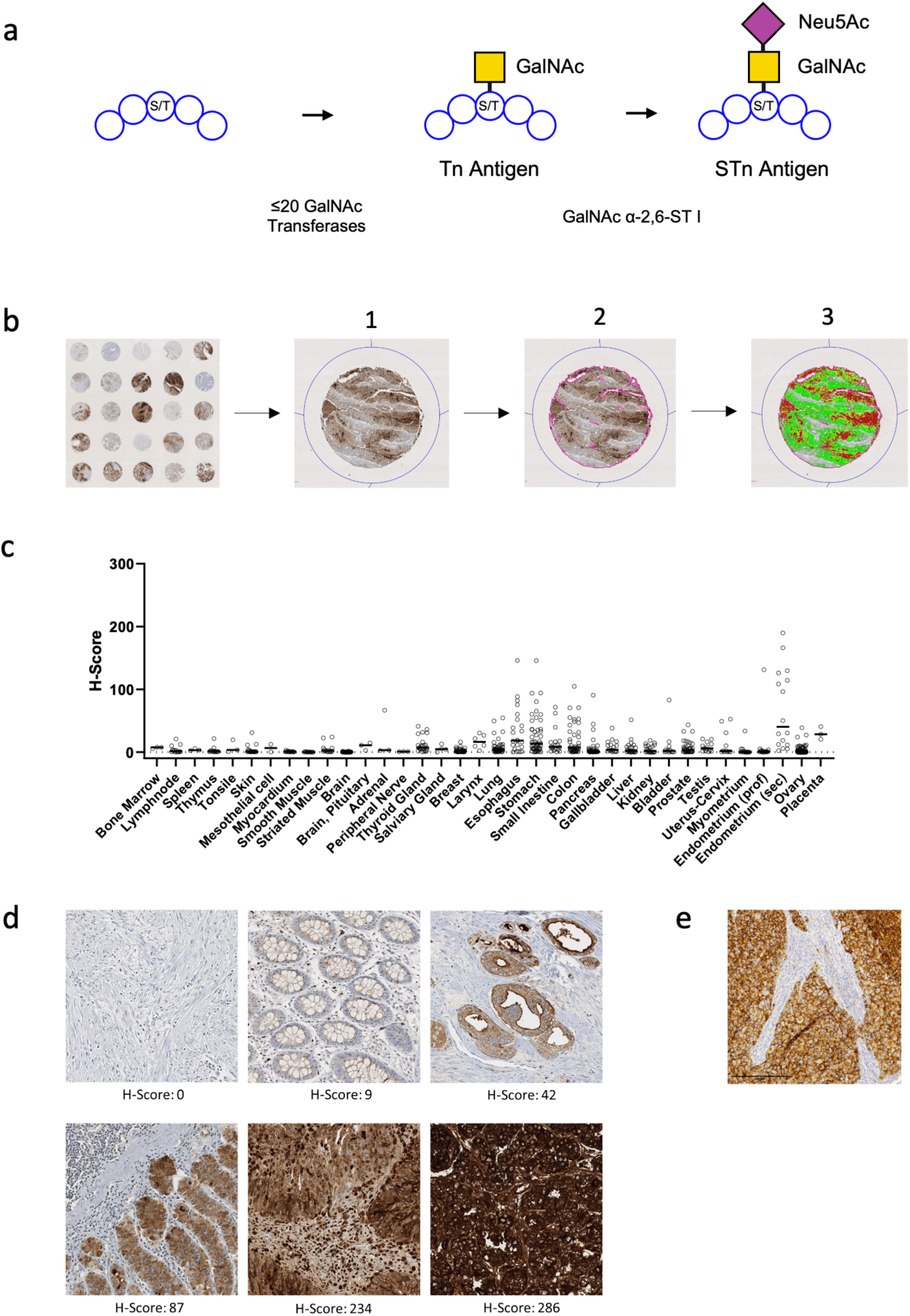
**a,** Synthesis pathway of Sialyl-Thomson-Nouveau (STn) antigen. Tn: Thomsen-Nouveau. S/T : serine / threonine. **b,** Processing of STn-stained tissue microarrays. 1:identifying cores, 2: identifying tissue, 3: partitioning into antigen-negative, low (0.2-0.4, green), medium (0.4-0.6, yellow) and strong (>0.6, red) areas by staining-intensity thresholds. **c,** H-score for STn expression of various healthy tissues. Black bar: median. **d,** Examples of anti-STn staining and corresponding H-score. **e,** STn-staining of A375-STn mouse tumor.

**Supplementary Data 2.**
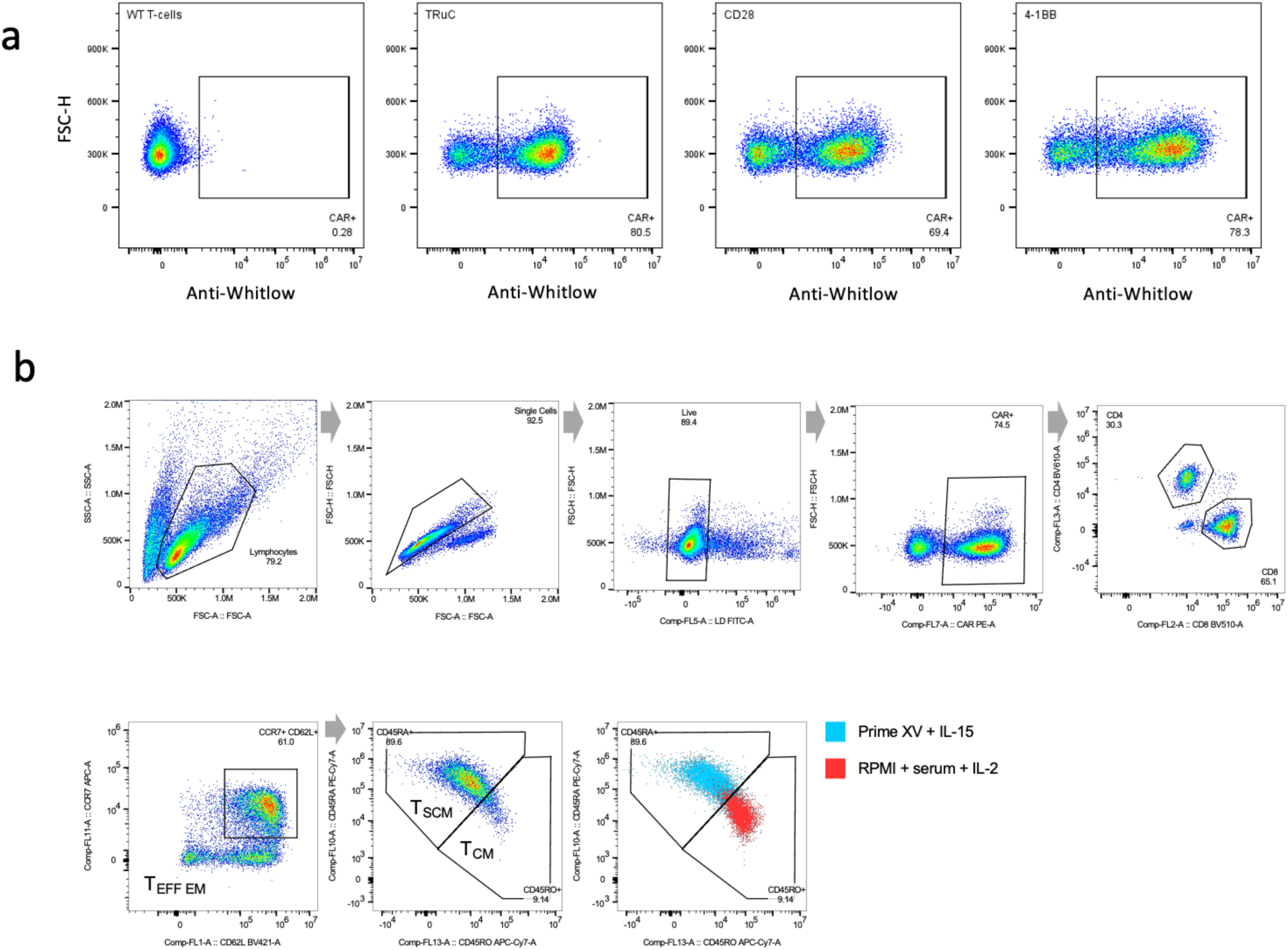
**a,** Staining of CAR-transduced T-cells with anti-Whitlow linker antibody. **b,** Gating strategy for phenotypic analysis. Example of CD45RA/CD45RO expression of T-cells cultured with different media.

**Supplementary Data 3.**
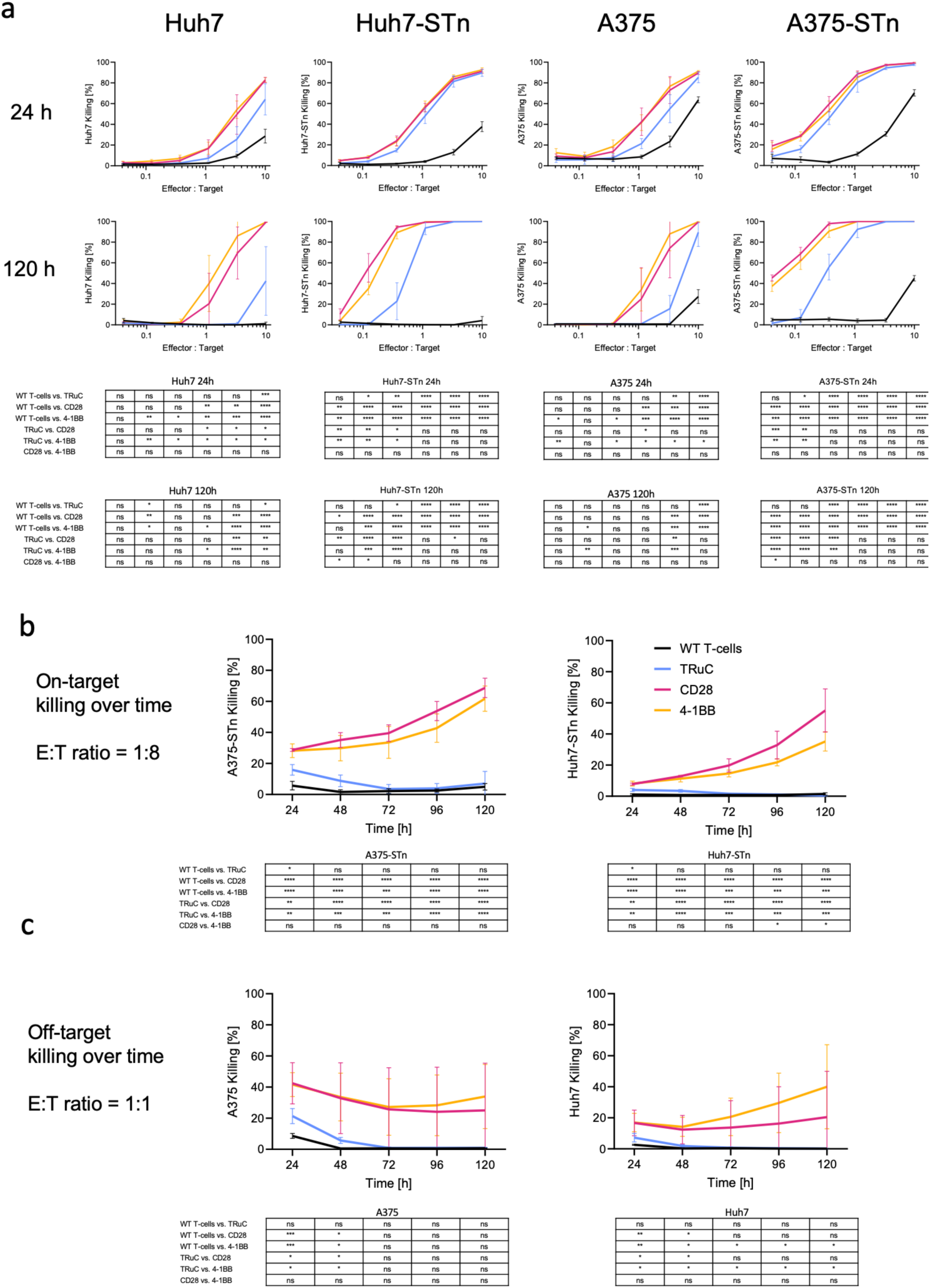
**a,** Cytotoxic activity of CAR-T cells and unmodified T-cells at different effector:target ratios at 24h and 120h of co-culture for STn-negative and positive cell lines. Pooled data of 4 different donors. Mean with STD shown. One-way ANOVA analysis of matched data for each time point or E:T ratio. **b,** Cytotoxic activity of CAR-T cells and untransduced T-cell over time against STn-positive A375 and Huh7 cancer cell lines. Pooled data of 4 different donors. Mean with STD shown. **c,** Cytotoxic activity of CAR-T cells and untransduced T-cell over time against STn-negative A375 and Huh7 cancer cell lines. Pooled data of 4 different donors. Mean with STD shown.

**Supplementary Data 4.**
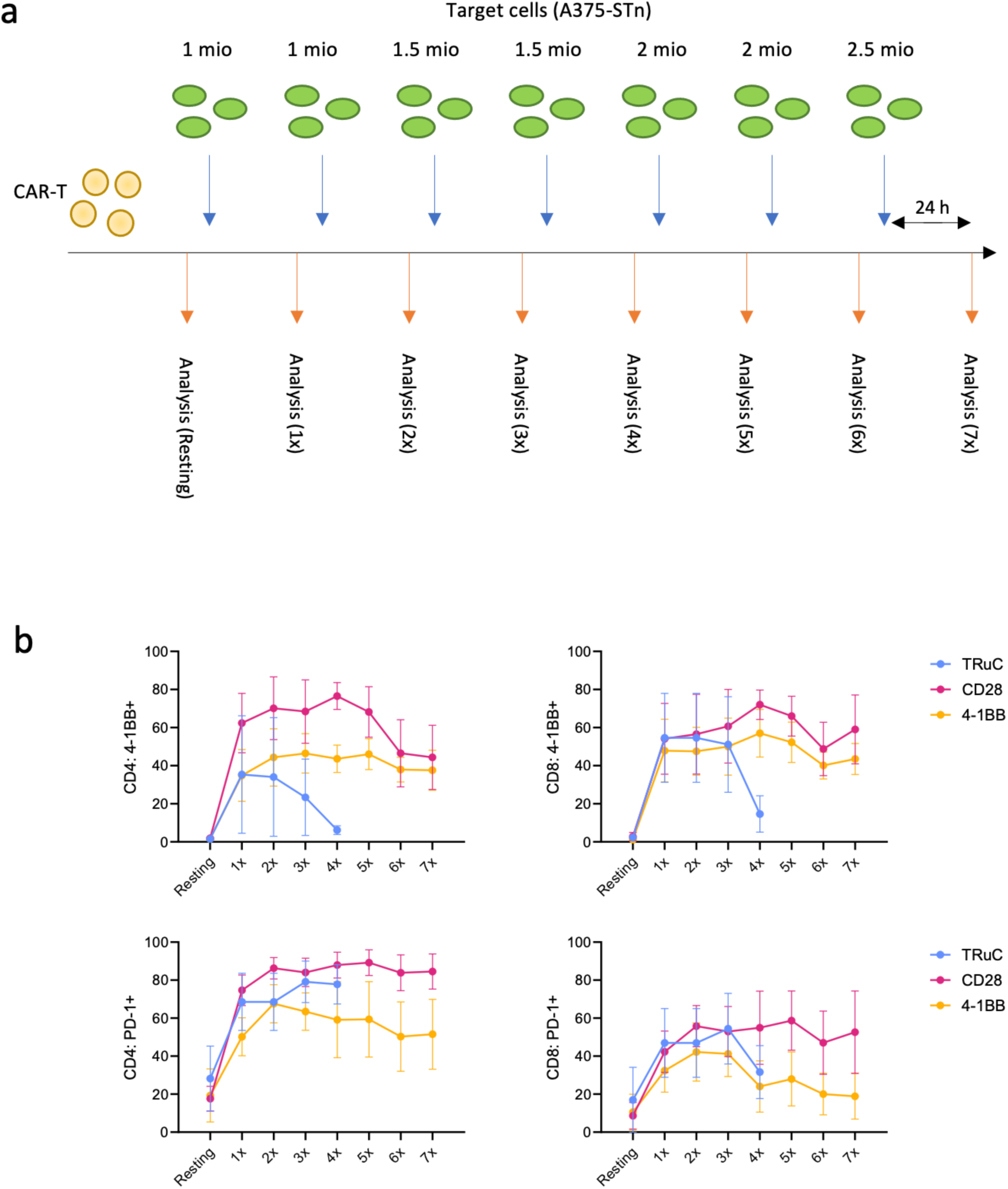
**a,** Experimental outline of repetitive stimulation experiment. CAR-T cells were stimulated twice a week with increasing number of target cells. **b,** PD-1 and 4-1BB expression for CD4 and CD8 CAR-T cells during repetitive stimulation after 24 hours of last stimulation. Pooled data of 4 different donors. Mean with STD shown.

**Supplementary Data 5.**
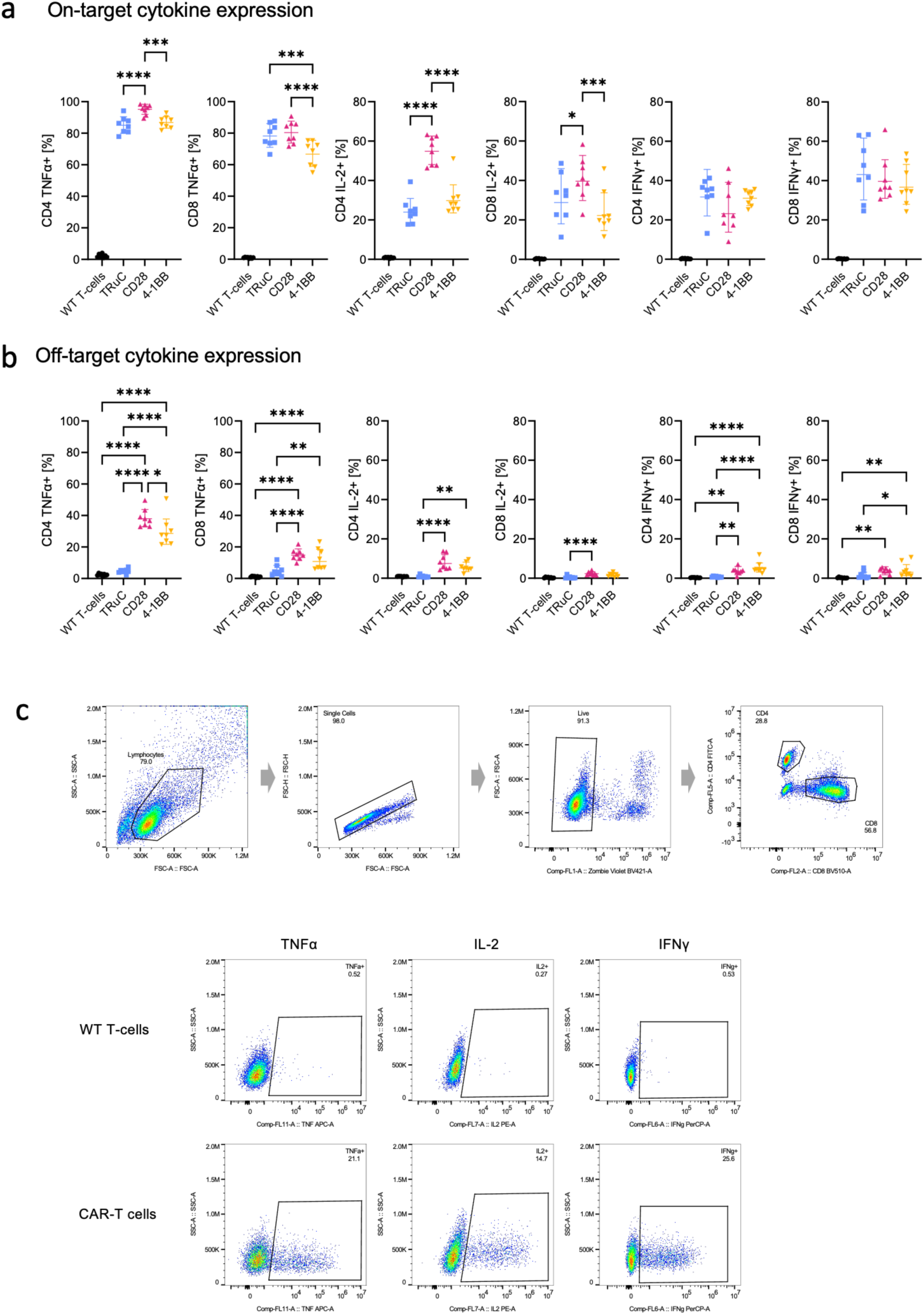
**a,** On-target intracellular cytokine expression of CAR-T cells when co-cultured with A375-STn cells. Statistics shown only for CAR variants. **b,** Off-target intracellular cytokine expression of CAR-T cells when co-cultured with A375 cells. Statistics shown only for CAR variants. **c,** Gating strategy for intracellular cytokine staining and comparison between anti-STn CAR-T cells and untransduced (WT: wildtype) T-cells.

**Supplementary Data 6.**
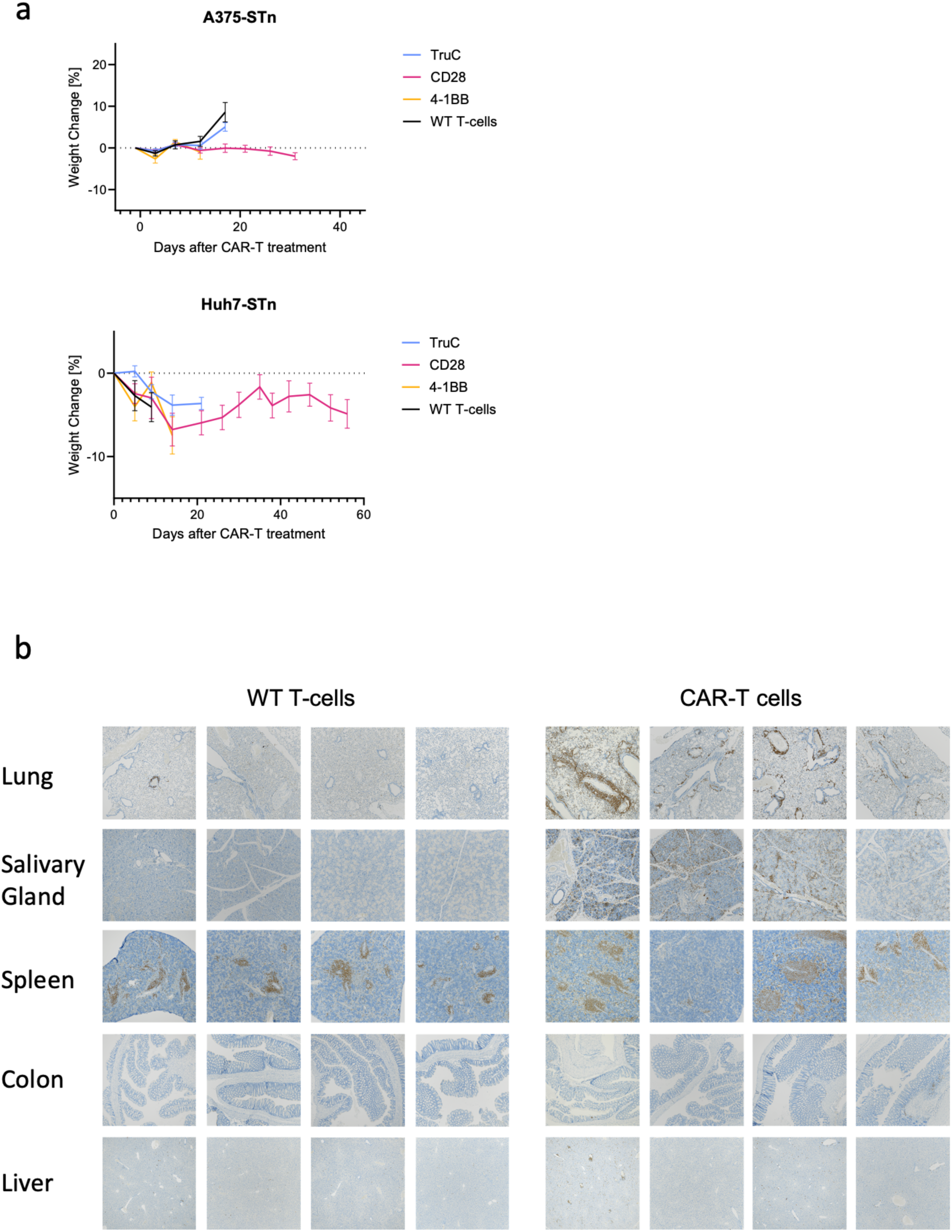
**a,** Weight change of animals after administration of 5 mio (A375-STn) or 15 mio (Huh7-STn) CAR-T cells. Data shown for time points where at least half of the animals in the treatment groups were alive. **b,** Biodistribution of CD3 cells within selected tissues for CAR and unmodified T-cells.

**Supplementary Data 7.**
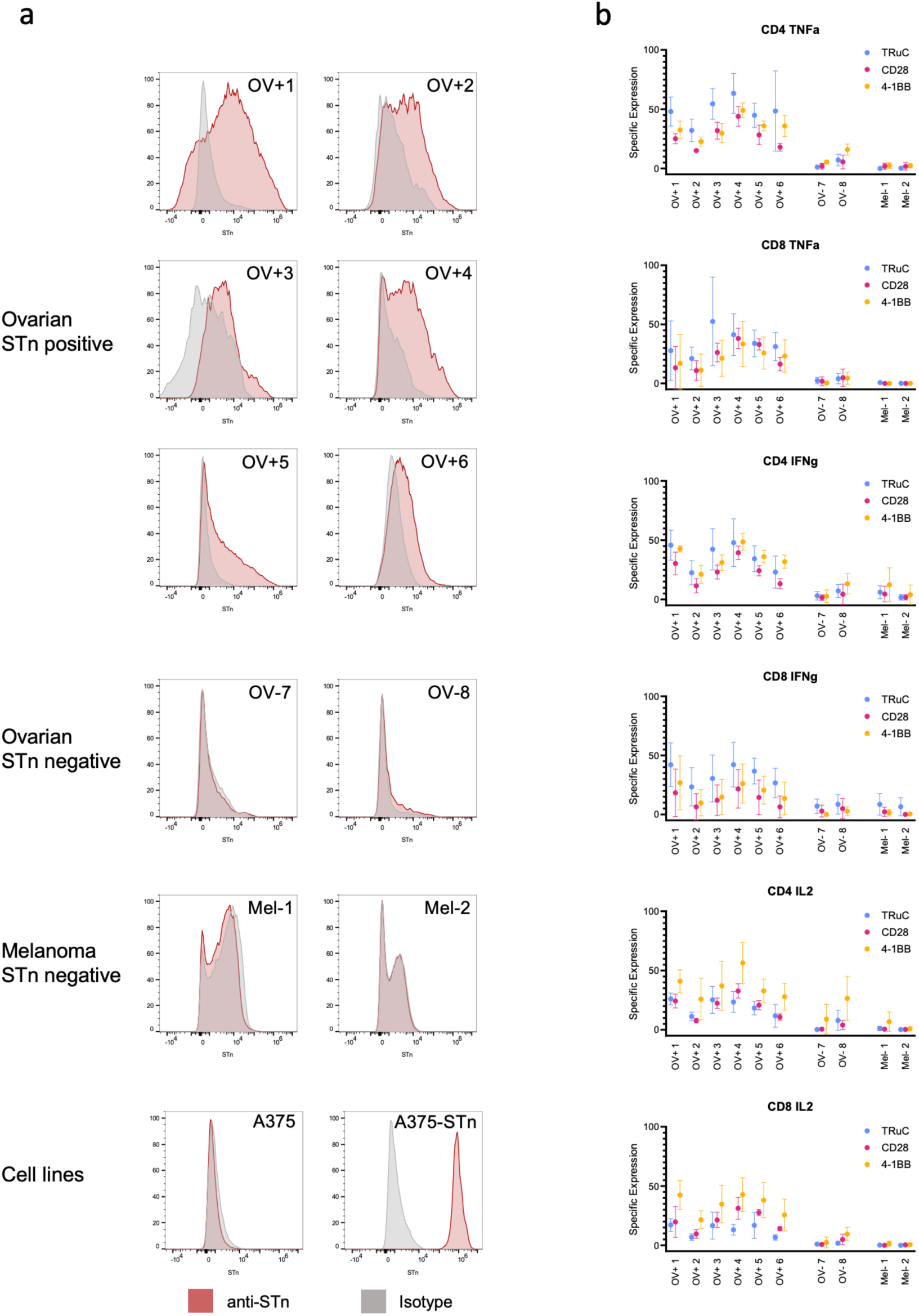
**a,** Anti-STn staining of primary tumor samples and cell lines used for co-culture assay. **b,** Intracellular cytokine staining of CD4 and CD8 CAR-T cells when co-cultured with primary tumor samples.

